# Easy Hi-C: A simple efficient protocol for 3D genome mapping in small cell populations

**DOI:** 10.1101/245688

**Authors:** Leina Lu, Xiaoxiao Liu, Jun Peng, Yan Li, Fulai Jin

**Affiliations:** Department of Genetics and Genome Sciences, Case Western Reserve University, 10900 Euclid Avenue, Cleveland, Ohio 44106, USA; Case Comprehensive Cancer Center, Case Western Reserve University, 10900 Euclid Avenue, Cleveland, Ohio 44106, USA

## Abstract

Despite the growing interest in studying the mammalian genome organization, it is still challenging to map the DNA contacts genome-wide. Here we present easy Hi-C (eHi-C), a highly efficient method for unbiased mapping of 3D genome architecture. The eHi-C protocol only involves a series of enzymatic reactions and maximizes the recovery of DNA products from proximity ligation. We show that eHi-C can be performed with 0.1 million cells and yields high quality libraries comparable to Hi-C.

---

The invention of Hi-C has transformed our understanding of the mammalian genome organizationx^1,2^. In the past few years, with increasing sequencing depth, a hierarchy of genome structures at different length scales has been revealed. With ∼10 million Hi-C reads at 1Mb resolution, it was first discovered that human genome is organized into two compartments (compartment A/B) reflecting the spatial separation of euchromatin and heterochromatin^1^. Later, with a few hundred million reads and at ∼50kb resolution, it was discovered that the mammalian genome is partitioned into Mb-sized topological associated domains (TADs), which are largely invariant between different cell types, and even conserved between mammalian species^3–7^. Most recently, kilobase resolution Hi-C analysis was achieved with sequencing depth at billion level^8,9^. At this resolution, cell type-specific chromatin loops within TADs between cisregulatory elements^8,10–13^, including promoters, enhancers, silencers and insulators, can be discerned. Apparently, with continuous improvement of sequencing technology, high-depth Hi-C data will become preferable because they can reveal genome organization with full detail.

However, generating high quality Hi-C library for deep sequencing is often challenging, especially when the amount of starting material is small. In Hi-C protocol, 5’ overhangs are created after restrictive DNA digestion (*e.g*. with *HindIII*) so that ligation junctions can be labeled with biotinylated nucleotides and eventually enriched in a pull-down step with streptavidin beads. However, this biotin-dependent strategy has several intrinsic limitations that affect library quality: (a) The efficiency of biotin incorporation into DNA is only ∼20-30%, sometimes as low as 5%^14^, therefore a majority of ligation junctions are not labeled; (b) Only a portion of labeled ligation junction products can be recovered after several washes, further decreasing the library complexity; (c) Biotin-pulldown may not completely remove contamination from unligated DNA products. Due to these reasons, Hi-C library from ∼10 million cells usually start to reach saturation after yielding several hundred million reads, and multiple biological replicates are necessary for kilobase-resolution Hi-C analysis. Therefore, most published Hi-C analysis are performed in cells or tissues where sample abundance is not a limiting factor.

We reasoned that we may circumvent the limitations of Hi-C, thus improve the assay efficiency by using a biotin-free strategy to enrich ligation products. Inspired by the well-established 4C method^15^, we developed easy Hi-C, which only involves a series of enzymatic reactions to generate DNA libraries for the study of genome architecture (**Fig. 1a**). In this new protocol, we begin with the *in situ* proximity ligation procedure and perform *HindIII* digestion and proximity ligation while keeping nuclei intact. It has been shown that comparing to “in-solution ligation” (ligation after nuclear lysis), the *in situ* ligation approach can reduce false positive interactions^9, 16, 17^. In eHi-C, ligation was performed directly after *HindIII* digestion without end repair, leading to *HindIII*-digestible junction products. After nuclear lysis and reverse crosslink, the DNA are digested with more frequent 4-base cutter *DpnII* before self-ligation (**Fig. 1a**). The samples are next treated with exonuclease to remove DNA that failed to form self-circles, and contaminations from un-ligated ends and other linear DNA species (**Fig. 1a**). In order to enrich the ligation junction products, we then employ a simple trick by cutting the self-circularized DNA again with *HindIII*, and only re-linearized junction DNA will be amplified during the library generation (**Fig. 1a**).

**Figure 1.**
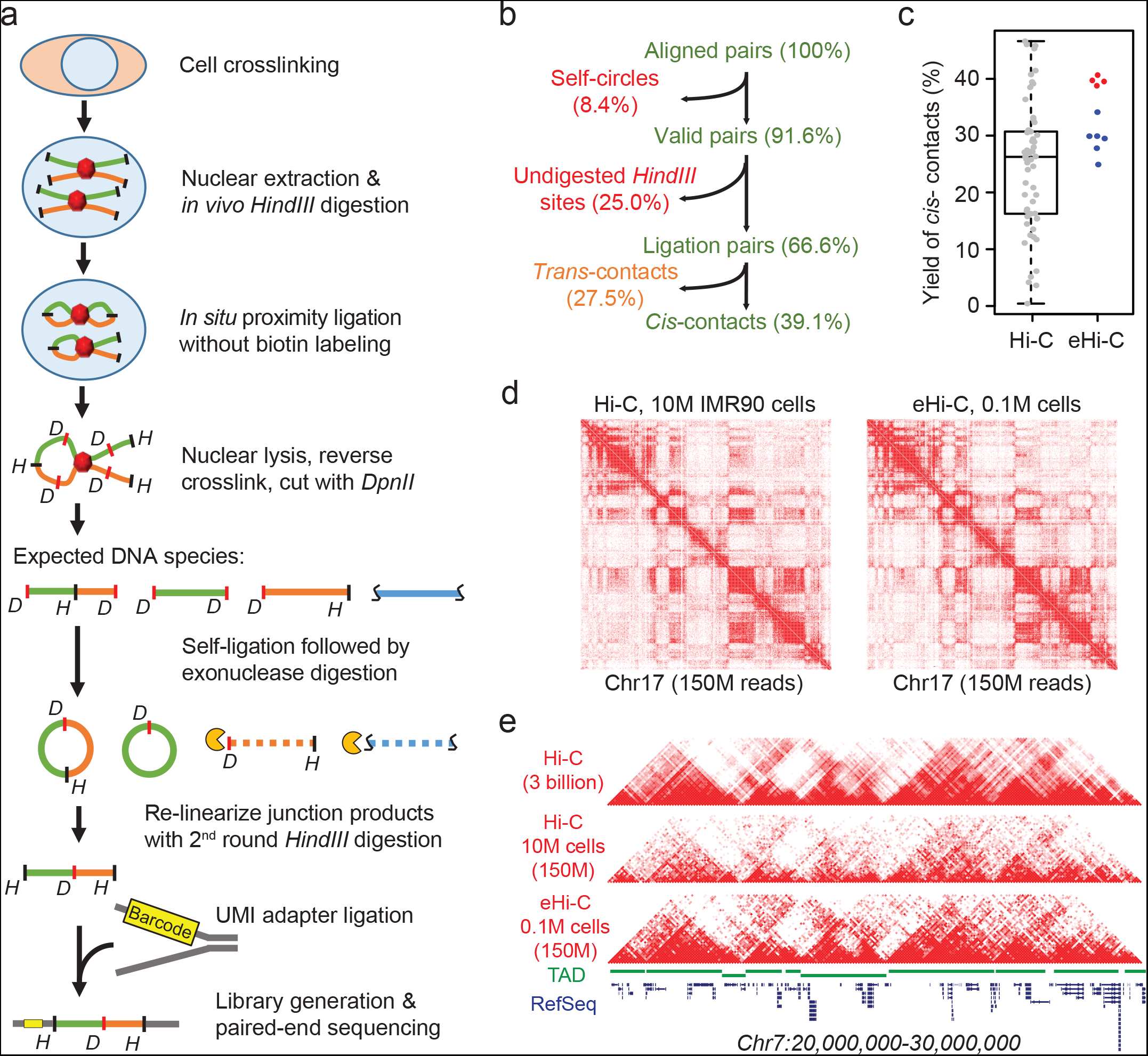
Easy Hi-C map DNA contacts genome-wide. The scheme of easy Hi-C. **b**, Data filtering results of one exemplary eHi-C library generated from 0.1M cells. **c**, Compare the yield of *cis*-contact reads between 57 published Hi-C libraries and 10 eHi-C libraries. The 4 red spots are eHi-C libraries prepared under *in situ* ligation condition. **d**, Heatmaps of contact matrices (chr17) from Hi-C and eHi-C at 200kb resolution. **e**, Heatmaps of contact matrices from Hi-C and eHi-C at 50kb resolution. The top track is drawn using a published IMR90 Hi-C dataset with ∼3 billion reads^8^. A track of TAD structures is plotted in green.

Because all the reads from eHi-C library are next to *HindIII* sites (**Fig. 1a**), the reproducible ligation events between the same pair of *HindIII* ends will give rise to identical paired-end reads (**Supplementary Fig. 1a**). This can be problematic because read pairs from the reproducible ligation events cannot be distinguished from PCR duplications. To address this issue, we used a custom adapter with a random sequence as unique molecule index (UMI) (**Fig. 1a**, **Supplementary Fig. 1b**). Two identical paired-end reads are considered PCR duplicates only if they have the same UMI sequences (**Supplementary Fig. 1b**). It should be also noted that a similar approach named ELP was developed several years ago to identify DNA contacts in fission yeast^18^. However, the design of ELP was flawed and cannot effectively remove contaminations from several species of non-junction DNA (**Supplementary Fig. 2**). Therefore, less than 4% of ELP reads represent proximity ligation events^18^. As mentioned above, by combining the *in situ* ligation procedure, an exonuclease treatment step, and the usage of UMI adapter (**Fig. 1a**), the eHi-C approach has solved the quality issues of ELP, and is suitable for the analysis of mammalian 3D genome.

We next tested the application of eHi-C in low-input setting with human primary lung fibroblast IMR90 cells. The eHi-C method uses more efficient “sticky end” ligation, and the protocol should havve a high recovery rate of ligation junction products because there is no DNA loss during the whole procedure (**Fig. 1a**). The only exception is the exonuclease digestion step: Ligation junction products may be digested if they fail to self-ligate (**Fig. 1a**). From a control experiment, we have determined that the efficiency of the self-ligation reaction is high (∼60%, **Supplementary Fig. 3**), thus a majority of ligation products can be recovered for sequencing. We have generated multiple eHi-C libraries each with 0.1∼0.2 million cells, and used low-depth or high-depth sequencing to evaluate the library quality (**Supplementary Table 1**). As expected, over 95% of mapped eHi-C reads begin with *HindIII* restrictive sequence AGCTT, indicating that nearly all reads are from re-linearized *HindIII*-digestible DNA circles. When one eHi-C library from 0.1 million cells is deep-sequenced to 150 million mapped read pairs, the percentage of PCR duplicates is low than all the 13 published IMR90 Hi-C libraries prepared with 100 times more (10 million) cells^8^ (**Supplementary Table 1–2**), indicating a significantly improved library complexity over Hi-C.

We next compared the sources of errors in Hi-C and eHi-C libraries by examining the fractions of different types of paired-end reads^8, 14^ (see **Method**, **Supplementary Fig. 4**). In Hi-C, read pairs fall in the same *HindIII* fragments are considered invalid, and the major type of invalid reads are “dangling reads” originated from non-ligation DNA (**Supplementary Fig. 4, Supplementary Table 2**). In contrast, the only type of invalid pairs from eHi-C are self-circles (**Supplementary Fig. 4b**), all the other types of invalid pairs are removed by exonuclease treatment. One drawback of eHi-C is that the data contain a significant number of false reads from undigested *HindIII* sites, which can be computationally removed as back-to-back read pairs next to the same restrictive sites (**Supplementary Fig. 4c**). We also filtered out *trans*- reads (two ends map to different chromosomes) because a big proportion of *trans-* reads often reflects high rate of random ligations^9, 14, 19^. After data filtering, we found that the yield of *cis*-contacts from eHi-C libraries, especially the ones prepared with *in situ* ligation procedure, is better than most of the published Hi-C libraries prepared with *HindIII*-based protocol (**Fig. 1c, Supplementary Table 1-2**). Importantly, the contact heatmaps from Hi-C and eHi-C data are identical showing the same component A/B^1^ and TAD^3^ structures (**Fig. 1d-e**). All these results demonstrated that eHi-C is a reliable alternative to Hi-C, and can correctly identify 3D genome features from small cell populations.

We also analyzed the intrinsic biases that may affect the eHi-C experimental procedure. Firstly, both Hi-C and eHi-C show a decay of contact frequency with increasing distance, but eHi-C captures more short range contacts in 1Mb range, presumably within TADs (**Fig. 2a**). Secondly, the contact frequencies involving very short *HindIII* restriction fragments (< 200bp) are lower in both Hi-C and eHi-C libraries, which can be explained by the spatial hindrance for those fragments to ligate^20^ (**Fig. 2b**). Interestingly, although eHi-C appears to preferentially captures contacts between mid-sized fragments (250∼1000bp), it has an overall alleviated bias compared to Hi-C, possibly because eHi-C uses more efficient sticky-end ligation instead of blunt-end ligation (**Fig. 2b**). Thirdly, we also observed GC-bias in eHi-C library, but it is intriguing that the profile of enrichment/depletion is opposite to what was observed for Hi-C^20^(**Fig. 2c**). We speculate that this might be because both ends of the eHi-C library starting from fixed *HindIII* restrictive sequence (AGCTT): the GC-bias in eHi-C reflects the efficiency of DNA polymerase elongation after it has already gone through first few bases during PCR amplification or sequencing. Finally, as expected, the length of ligation product will also affect the efficiency of DNA self-circularization (**Fig. 2d**). These results provide a basis for the eHi-C data normalization and computational inference of DNA contacts.

**Figure 2.**
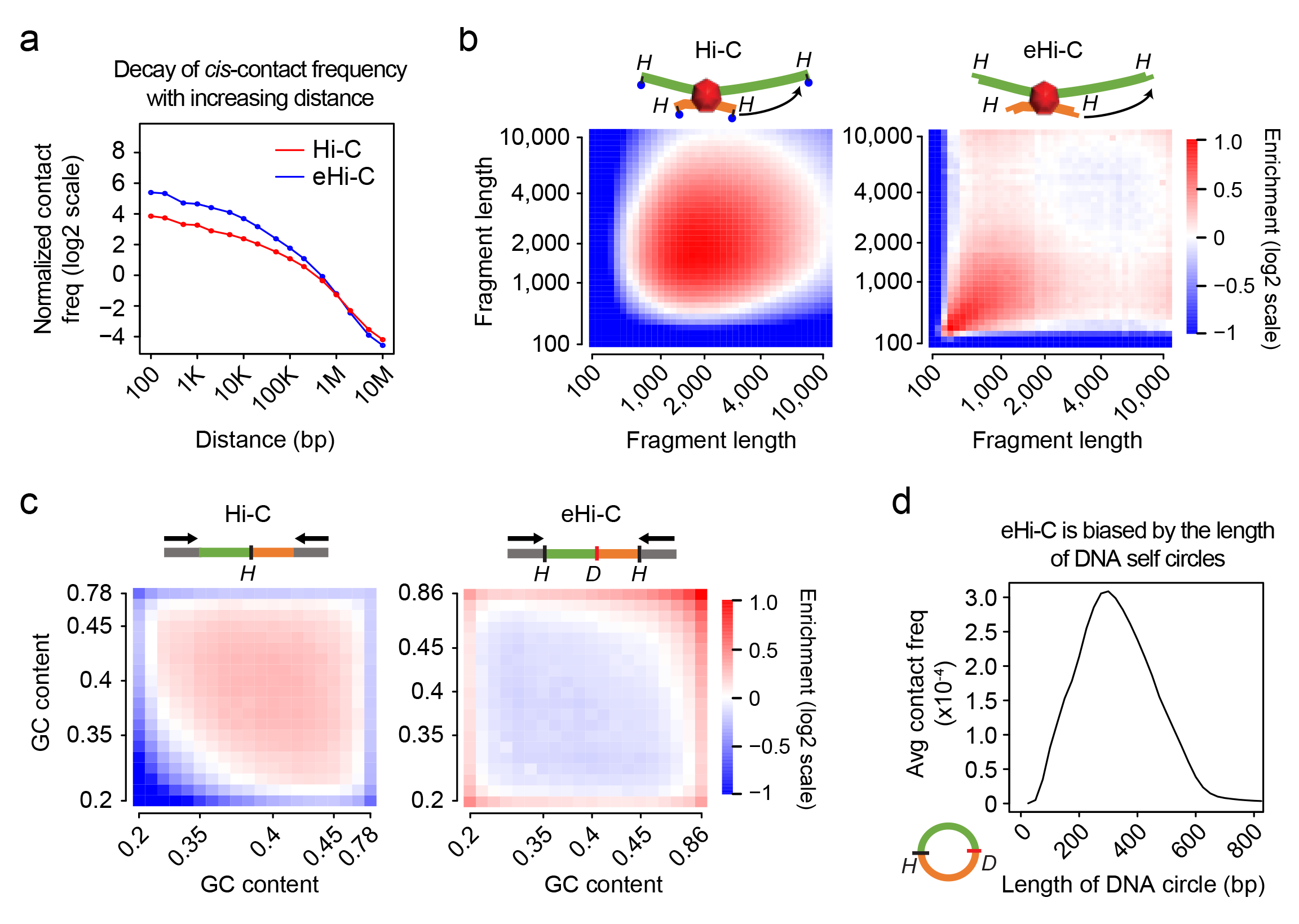
Systematical biases in eHi-C experiments. **a**, Curves showing the decay of *cis*-contact with an increasing distance between two *HindIII* restrictive fragments. Only “same-strand” reads (see **Methods**) were used to plot the curves. **b**, Compare the bias from *HindIII* fragment length in Hi-C (left) and eHiC (right) data. All the fragments are binned into 40 equal-sized groups, and the enrichment of *trans* reads between any two groups are plotted as heatmaps. The enrichment value is defined by the ratio between actual read counts and the global average for any two groups. **c**, Similarly, *HindIII* ends are binned into 20 groups based on GC content, and the enrichment of *trans*- reads are also plotted as heatmaps. For Hi-C (left) we used the GC content in the 200bp region upstream the *HindIII* site, and for eHi-C, we used the GC content of the region between the *HindIII* and its nearest *DpnII* site. **d**, Curve shows the average contact frequency from eHi-C against the length of ligation junction products forming DNA circles.

To summarize, we have developed a novel easy Hi-C method for convenient mapping of 3D genome architecture. It relies on a highly efficient biotin-free strategy to enrich ligation junctions and therefore is capable of generating high quality DNA library from as few as 0.1 million cells. This low-input eHi-C approach will facilitate the study of genome architecture in rare tissues or small cell populations which are previously unachievable with conventional Hi-C technology.

## METHODS

### Cell culture and fixation

Human primary IMR90 fibroblasts were grown as previously described^8^. After confluence, the cells were detached with trypsin and collected by spinning down at 900g for 5 minutes. Then the cells were fixed in 1% formaldehyde for 15 minutes at 37ºC, followed by 1/20 volume of 2.5M glycine at room temperature for 5 minutes to quench formaldehyde. The fixed cells were scrapped, washed in PBS and pelleted before stored in -80°C.

### Easy Hi-C

In this study low-input eHi-C libraries were prepared in two settings. In the first scenario (“aliquot” setting), we started with 1 million IMR90 cells and go through the protocol described below and usually resulted in 250∼500ng DNA for library preparation (**Fig. 1a**). 10% or 20% of these DNA were used to generate library (0.1 or 0.2 million cells per library). In the second scenario (“mini” setting), we started the experiments with lysing 0.1 or 0.2 million cells following the same protocol as described below, except that all steps before library preparation were performed in 25% volume. Because the cell lysis and *HindIII* digestion conditions in this work are different from the published *in situ* Hi-C protocol. We have made modifications in order to ensure nuclei integrity during ligation.

#### Cell lysis, HindIII digestion, and in situ ligation

Cell pellet from 1 million cells was lysed in 1ml cell lysis buffer (10mM Tris-Cl, pH7.5, 10mM NaCl, 0.2% NP-40, 1X proteinase inhibitor cocktail (Roche)) before incubating on ice for 15 minutes. If there is cell clump in the tube, we dounce the cells for 10 times every cycle for 3 cycles, with one-minute on ice between each cycle. After douncing, the nuclei were put on ice for another 5 minutes and then pelleted by centrifuging (5,000rpm for 5 minutes at 4°C). The pellets were washed once in 1X Cutsmart buffer (NEB) before resuspended in 360ul 1× Cutsmart buffer. After resuspension, 40ul of 1% SDS were added (final 0.1%), and the tubes were incubated at 65ºC for 10 minutes. To quench the SDS, 44ul of 10% Triton X-100 (final 1%) was then added to each tube. For chromatin digestion, 400U *HindIII* (NEB, R3104M 100U/µl) were added to each tube followed by incubation at 37°C for 4 hours. To ensure efficient digestion, another 400U of *HindIII* were added to each tube again for overnight digestion. On day 2, we digested the nuclei for another 4 hours by adding fresh *HindIII* enzyme (400U). After digestion, the enzyme was inactivated by adding 40ul of 10% SDS (final 1%) to each tube and incubation at 65°C for 20 minutes. The digested products were then transferred to a new 15ml tube and mixed with 3.06ml 1.15X ligation buffer (75.9mM Tris-HCl, ph7.5, 5.75mM DTT, 5.75mM MgCl_2_ and 1.15mM ATP). 187ul 20% Triton X-100 was added to the mixture and incubated at 37ºC for 1 hour. For ligation, the products were then mixed with 30ul of T4 DNA ligase (Invitrogen, 15224-025, 1U/ul) and incubated at 16ºC overnight. After ligation, the tubes were put in room temperature for 30 minutes and the nuclei were pelleted by centrifuging at 5,000rpm for 5 minutes. The supernatant was discarded to remove the free DNA and only the nuclei pellets were kept. The nuclei pellet step is skipped in the “dilute” libraries in **Supplementary Table 1**. The nuclei pellets were then resuspended in 3.06ml of 1.15X ligation buffer and mixed with 40ul of 10% SDS and 187ul of 20% Triton X-100 for nuclear lysis.

#### Reverse crosslinking, DpnII digestion and self-ligation

After nuclear lysis, the mixture were then reverse crosslined at 65°C overnight after adding 25ul of 20mg/ml proteinase K. DNA were purified with Phenol: Chloroform: Isoamyl Alcohol (25:24:1, Sigma) following standard protocol. 2∼3μg DNA are expected from 1M cells. The DNA was then digested with 50U *DpnII* (NEB, R0543L, 10U/μL) in a total volume of 100uL at 37°C for 2 hours. After digestion, the enzyme was heat inactivated at 65°C for 25 minutes. The mixture was first incubated with 0.5 volume of PCRClean DX beads (Aline Biosciences) at room temperature for 10 minutes before harvest the supernatant according to vendor’s protocol. The supernatant was then incubated with 2 volume of PCRClean DX beads at room temperature for 10 minutes. DNA on the beads were then harvested in 300ul nuclease free water. The two-step bead purification results in DNA with a size range 100 ∼ 1,000bp. The DNA products were then mixed with 200ul of 5X ligation buffer, 5U T4 DNA ligase (Invitrogen) and water to a total volume of 1ml. Self-ligation was done by incubating the tubes at 16ºC overnight.

#### Exonuclease digestion and DNA circle re-linearization

The self-ligated DNA were purified again with Phenol: Chloroform: Isoamyl alcohol and digested with 6U of lambda exonuclease (NEB, M0262S) in 200μL volume at 37°C for 30 minutes. The exonuclease was then inactivated by incubating at 65°C for 20 minutes. Resulting DNA were purified with 2 volume of PCRClean DX beads as described above. For DNA circle re-linearization, bead bound DNA were eluted and digested with 20U *HindIII* again at 37°C for 2 hours in 150μL volume. The *HindIII* enzyme was inactivated at 65°C for 20 minutes, and the DNA was purified with 2 volume PCRClean DX beads for another time as described above. In the end, bead-bound DNA was eluted in 50ul nuclease free water. From 1M cells, we expect 250-500ng DNA in the end.

#### Library preparation

We took 10∼20% of re-linearized DNA (∼50ng) for library generation following Illumina TruSeq protocol. Briefly, the DNA was firstly end repaired using End-it kit (Epicentre). The end-repaired DNA was then A tailed with Klenow fragment (3′–5′ exo–; NEB) and purified with PCRClean DX beads. Bead bound DNA were eluted in 20μL water and then reduced to 4μL using Speedvac at 50°C. The 4ul DNA product was mixed with 5ul of 2× quick ligase buffer, 1ul of 1:10 diluted annealed adapter and 0.5ul of Quick DNA T4 ligase (NEB). The ligation was done by incubating at room temperature for 15 minutes and the enzyme was then inactivated by incubating at 65°C for 10 minutes. DNA was then purified with 1.8 volume of DX beads as described above. Elution was done in 14ul nuclease free water. When we check eHi-C libraries quality, we only need to sequence less than 1 million reads on MiSeq (Illumina). Because the proportion of PCR duplicates from low-depth sequencing is very low, we used TruSeq indexed adapters (Illumina) without UMI barcode. When we need to deep sequence an eHi-C library, we used custom TruSeq adapter in which the index is replace by 6 base random sequence. The custom adapter was generated by annealing the following two oligos:

Universal oligo –

AATGATACGGCGACCACCGAGATCTACACTCTTTCCCTACACGACGCTCTTCCGA TC*T

UMI oligo –

/5Phos/GATCGGAAGAGCACACGTCTGAACTCCAGTCACNNNNNNATCTCGTATG CCGTCTTCTGCTT*G

#### PCR amplification of DNA libraries

To amplify the DNA libraries, we mixed 13ul adapter ligated DNA with 1ul of 20uM oligo C (AATGATACGGCGACCACCGAGATCTACAC), 1ul of 20uM oligo D (CAAGCAGAAGACGGCATACGAGAT) and 15ul of 2X KAPA HiFi Hotstart ready mix (Kapa Biosystems). And the PCR amplification was done as follows: denature at 98ºC for 45 seconds, cycled at 98°C for 15 seconds, 60°C for 30 seconds, 72°C for 30 seconds, and did 5 cycles at first for estimating the total cycle number needed, and then further extension at 72°C for 5minutes. The products were then purified using 1.8 volume of DX beads to remove primer contamination as described above. And the DNA was eluted in 20ul nuclease free water. And library quantification was done following the protocol of Illumina library quantification kit (KAPA Biosystems, KK4824). PCR was done again in 50μL volume for a target final concentration 20∼40nM (usually 3∼4 additional cycles). The generated libraries were then subjected to sequencing.

### Analysis of Hi-C and eHi-C data

#### Alignment and removing PCR duplications

Published IMR90 Hi-C data were used in this study to compare with eHi-C. The accession numbers of Hi-C data are listed in **Supplementary Table 2**. All the sequencing data were mapped to human reference genome hg18 using Bowtie. For Hi-C, the two ends of paired-end (PE) reads were mapped independently using the first 36 bases of each read. PCR duplications were defined as PE reads with both ends mapped to the same locations. For eHi-C, because nearly all the mappable reads start with *HindIII* sequence AGCTT, we trimmed the first 5 bases from every read, took the next 36 bases, and add the 6-base sequence AAGCTT to the 5’ of each reads before mapping using the whole 42 bases. Some MiSeq runs were performed with reads shorter than 41 bases, and the full length reads will be used in those cases. After mapping, we further filter the reads requiring the positions of both ends to be exactly at the *HindIII* cutting sites. Because the eHi-C library deep sequenced was prepared with UMI adapter, PCR duplications were defined as identical PE reads also with the same UMI barcode. The eHi-C libraries sequenced on MiSeq were not intended for deep sequencing and therefore do not have UMI barcode. PCR duplications in those libraries were removed the same way as Hi-C.

#### QC analysis of Hi-C and eHi-C libraries

After remove PCR duplications, we analyzed the library quality by classifying the reads into different categories. In both Hi-C and eHi-C, the percentage of *trans*- contacts can be easily calculated by count the number of reads with two ends on different chromosomes (listed in **Supplementary Table 1-2**). For *cis*- reads in Hi-C data, we first discard the reads with both ends mapped to the same *HindIII* fragments as invalid pairs. Dangling ends are defined as “inward” pairs among the invalid pairs (**Supplementary Fig. 4a**) and the percentage are listed in **Supplementary Table 2**. The rest invalid pairs are classified into “other false” category. Since cut-and-ligation events are expected to generate reads within 500bp upstream of *HindIII* cutting sites due to the size selection (“+” strand reads should be within 500bp upstream of a *HindIII* site, and “+” strand reads should be within 500bp downstream a *HindIII* site), we only keep reads pairs with both ends satisfying this criteria. The other pairs are also classified into “other false” category in **Supplementary Table 2**. After this step, every remaining paired-end reads represents one pair of restrictive fragments. We next split all these reads into three classes based on their strand orientations (“same-strand”, “inward”, or “outward”). We have previously shown that although theoretically “same-strand” reads should be twice as many as “inward” or “outward” reads, in reality more “inward” or “outward” reads can be observed due to incomplete digestion of chromatin^8^. We therefore estimate the total number of real *cis*-contact as twice the number of valid “same-strand” pairs, the percentages are listed in **Supplementary Table 2**. For eHi-C, library, the only type of invalid cis- pairs are self-circles with two ends within the same *HindIII* fragment facing each other (**Supplementary Fig. 4b**). Similar to Hi-C, we also computed the total number of real *cis*-contact as twice the number of valid “same-strand” pairs. Reads from undigested *HindIII* sites are back-to-back read pairs next to the same *HindIII* sites facing away from each other (**Supplementary Fig. 4c**).

#### Bias analysis of Hi-C and eHi-C libraries

To plot the decay of contact with distance (**Fig. 2a**), we only used “same-strand” *cis*-contact reads. For any given distance *L*, we found all *HindIII* fragment pairs with gap distance between 0.9 ∗ *L* and 1.1 ∗ *L*, and computed the average contact frequency among them. We normalize these numbers by dividing them by the average contacts from all the intra-chromosome *HindIII* fragment pairs. We used *trans*- contact data to compute the fragment length and GC-content bias because in this way, the distance is no longer a parameter of concern.

For length bias (**Fig. 2b**), we divided all the *HindIII* fragments into 40 equal-sized groups and computed the average contact frequency for each pair of groups, and enrichment values were calculated by normalizing to global average. Similarly, we plot the GC bias (**Fig. 2c**) by dividing all the *HindIII* ends into 20 equal-sized groups by GC content. For Hi-C, the GC content was computed using the 200bp near each *HindIII* end. For eHi-C, the GC content was computed for the region between a *HindIII* end and the nearest *DpnII* site.

#### Data availability

Raw and processed eHi-C data are available at NCBI GEO with accession number GSExxxxx.

